# Undermatching is a consequence of policy compression

**DOI:** 10.1101/2022.05.25.493472

**Authors:** Bilal A. Bari, Samuel J. Gershman

**Affiliations:** Department of Psychiatry, Massachusetts General Hospital, Boston, MA 02114; Department of Psychology and Center for Brain Science, Harvard University, Cambridge, MA 02138; Center for Brains, Minds, and Machines, MIT, Cambridge, MA 02139

**Author notes:** Correspondence: Samuel J. Gershman.

## Abstract

The matching law describes the tendency of agents to match the ratio of choices allocated to the ratio of rewards received when choosing among multiple options (Herrnstein, 1961). Perfect matching, however, is infrequently observed. Instead, agents tend to undermatch, or bias choices towards the poorer option. Overmatching, or the tendency to bias choices towards the richer option, is rarely observed. Despite the ubiquity of undermatching, it has received an inadequate normative justification. Here, we assume agents not only seek to maximize reward, but also seek to minimize cognitive cost, which we formalize as policy complexity (the mutual information between actions and states of the environment). Policy complexity measures the extent to which an agent’s policy is state-dependent. Our theory states that capacity-constrained agents (i.e., agents that must compress their policies to reduce complexity), can only undermatch or perfectly match, but not overmatch, consistent with the empirical evidence. Moreover, we validate a novel prediction about which task conditions exaggerate undermatching. Finally, we argue that a reduction in undermatching with higher dopamine levels in patients with Parkinson’s disease is consistent with an increased policy complexity.

**Significance statement:** The matching law describes the tendency of agents to match the ratio of choices allocated to different options to the ratio of reward received. For example, if option A yields twice as much reward as option B, matching states that agents will choose option A twice as much. However, agents typically undermatch: they choose the poorer option more frequently than expected. Here, we assume that agents seek to simultaneously maximize reward and minimize the complexity of their action policies. We show that this theory explains when and why undermatching occurs. Neurally, we show that policy complexity, and by extension undermatching, is controlled by tonic dopamine, consistent with other evidence that dopamine plays an important role in cognitive resource allocation.

## Introduction

Over half a century ago, Richard Herrnstein discovered an orderly relationship between choices and rewards (Herrnstein, 1961), which he termed ‘matching’ behavior. Herrnstein’s matching law describes the tendency of animals to ‘match’ the ratio of choices allocated to the ratio of reward received when choosing among multiple options. For 2 options, matching is defined by:

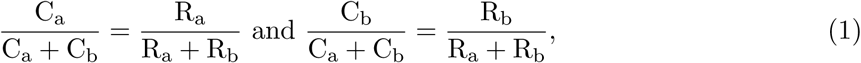

where C_a_ is the number of choices allocated to option a and R_a_ is the number of rewards obtained from option a. The matching law describes choice behavior fairly accurately in a number of animals, including pigeons (Herrnstein, 1961; de Villiers and Herrnstein, 1976; Baum, 1979; Mazur, 1981; Villarreal et al., 2019), mice (Gallistel et al., 2007; Fonseca et al., 2015; Bari et al., 2019), rats (Graft et al., 1977; Gallistel, 1994; Belke and Belliveau, 2001; Gallistel et al., 2001; Lee et al., 2017), monkeys (Anderson et al., 2002; Sugrue et al., 2004; Lau and Glimcher, 2005; Kubanek and Snyder, 2015; Tsutsui et al., 2016; Soltani et al., 2021), and humans (Schroeder and Holland, 1969; Pierce and Epling, 1983; Beardsley and McDowell, 1992; Savastano and Fantino, 1994; Vullings and Madelain, 2018; Cero and Falligant, 2020).

Perfect matching, however, is only seen infrequently. Theoretically, animals can deviate from perfect matching by undermatching, biasing choices towards the poorer option, or by overmatching, biasing choices towards the richer option (Baum, 1974). Empirically, animals systematically *undermatch* (Baum, 1974; Myers and Myers, 1977; Baum, 1979; Wearden and Burgess, 1982). Overmatching is rarely observed. This systematic bias towards undermatching has received a number of explanations, including mistuned learning rules (Loewenstein and Seung, 2006; Loewenstein, 2008), procedural variation in tasks (Baum, 1979; Williams, 1985), the inability to detect the richer stimulus/action (Baum, 1974), poor credit assignment (Trepka et al., 2021), inappropriate choice stochasticity (Trepka et al., 2021), noise in neural mechanisms of decision making (Soltani et al., 2006), synaptic plasticity rules (Iigaya and Fusi, 2013), belief in environmental volatility (Saito et al., 2014), and optimal decision-making under uncertainty (Iigaya et al., 2019). None of these models explain the conspicuous absence of overmatching in the empirical literature. Here, we provide a normative rationale for undermatching and the relative absence of overmatching.

We begin with the premise that agents seek to simultaneously maximize reward and minimize some measure of cognitive cost, which we formalize as *policy complexity*, the mutual information between states and actions (Parush et al., 2011; Gershman, 2020; Lai and Gershman, 2021). The policy *π*(*a*|*s*) corresponds to the probabilistic mapping between environment states (*s*) and actions (*a*). Because policy complexity is a lower bound on the number of bits needed to store a policy in memory, more complex policies necessitate more bits. If the optimal policy exceeds an agent’s memory capacity, then it will need to “compress” its policy by reducing complexity. In this paper, we argue that undermatching is a consequence of policy compression.

We first extend the notion of policy complexity to describe matching behavior. We find that agents should only perfectly match or undermatch, but never overmatch, since overmatching requires more memory than perfect matching and yields less reward. We validate a novel prediction that capacity-constrained agents should increase undermatching on task variants that demand greater policy complexity for perfect matching behavior. We then test an implication of the hypothesis that tonic dopamine signals average reward (Niv et al., 2007), and thereby controls the allocation of cognitive effort (Mikhael et al., 2021). When tonic dopamine is higher, individuals should adopt higher policy complexity, as if their capacity limit had effectively increased. We find evidence for this hypothesis using data from patients with Parkinson’s disease performing a dynamic foraging task on and off dopaminergic medication (Rutledge et al., 2009): undermatching was reduced on medication compared to off medication. Taken together, our results support a policy compression account of undermatching.

## Materials and Methods

### Behavioral data

We reanalyzed data from mice (Bari et al., 2019) and human subjects (Rutledge et al., 2009) performing a dynamic foraging task (differences between the mouse and human versions of the task are detailed below). In this task, subjects chose between two options, each of which delivered reward probabilistically. The ‘base’ reward probabilities of each option remained fixed within a given block, and changed between blocks. Block transitions were uncued. A key feature of the dynamic foraging task was the baiting rule, which stipulated that the longer an agent has abstained from choosing a particular option, the greater the probability of reward on that option. Stated another way, if the unchosen option would have been rewarded, the reward was delivered the next time that option was chosen. The baiting rule was designed to mimic ecological conditions, where abstaining from a foraging option will allow that option to replenish reward. Mathematically, the baiting probability took the following form:

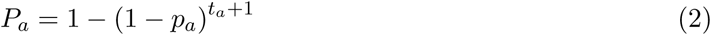

where *p*_*a*_ is the base reward probability for option *a, t*_*a*_ is the number of consecutive choices since that option was last chosen, and *P*_*a*_ is the probability of reward when the agent next chooses that option (illustrated in Figure 1A).

**Figure 1:**
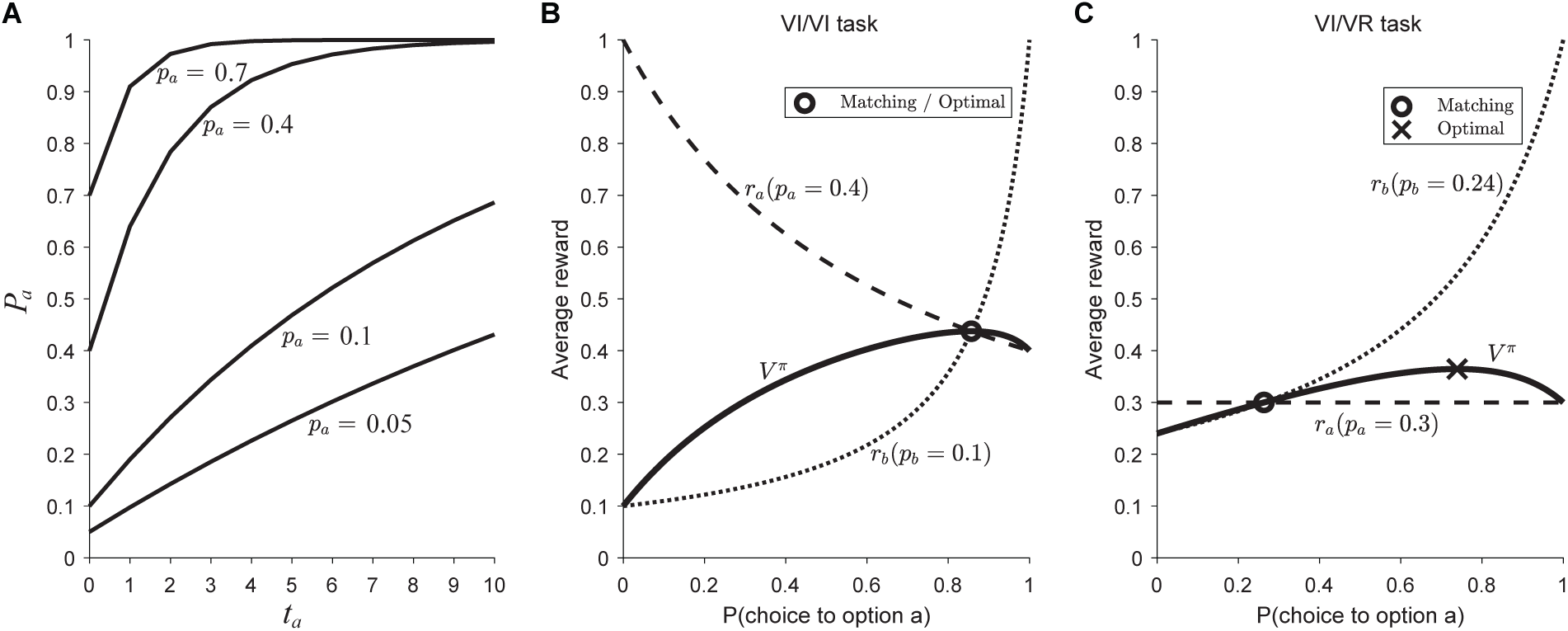
Geometric representation of matching behavior in different task conditions. A: The baiting rule for four different values of *p*_*a*_ (base reward probabilities). The x-axis is *t*_*a*_, the number of consecutive choices since that option was last chosen, and the y-axis is *P*_*a*_, the probability of reward. Here, *p*_*a*_ ∈ {0.05, 0.1, 0.4, 0.7}. Adapted from Huh et al. (2009). B: In variable-interval / variable-interval (VI/VI) tasks, where both options employ the ‘baiting’ rule, matching emerges as the optimal probabilistic policy. Matching occurs where *r*_*a*_ = *r*_*b*_ (hence they ‘match’). C: Variable-interval / variable-ratio (VI/VR) tasks allow us to disambiguate whether animals match or whether they approximate the optimal probabilistic solution. In these task variants, one option (here, option *b*) follows a VI schedule (i.e., programmed with the baiting rule) and the other option (here, option *a*) follows a VR schedule (standard probabilistic reward delivery). The matching policy (where *r*_*a*_ = *r*_*b*_) differs from the optimal probabilistic policy. Adapted from Bari and Cohen (2021).

In the mouse dynamic foraging task, male C57BL/6J mice (The Jackson Laboratory, 000664), ages 6-20 weeks, were surgically implanted with a head plate in preparation for head fixation. After recovery, these animals were water restricted and habituated. Cues were delivered in the form of odors via a custom-made olfactometer (Cohen et al., 2012). Animals received one of two cues. The “go cue”, delivered on 95% of trials, signaled to the animal to make a leftward or rightward lick towards a custom-built dual lick port. The neutral “no-go cue”, delivered on 5% of trials, was neither rewarded nor punished, and we ignore all no-go trials for analyses in this paper. We included data from two task variants. In the “40/10” task variant, the base reward probabilities switched between {0.4/0.1} (corresponding to base reward probabilities for the left and right option) and {0.1/0.4}. This corresponds to a task with two possible states. A similar logic applied to the “40/5” task variant. We included 23 mice total, 10 of which performed the 40/10 task for 236 total sessions and 17 of which performed the 40/5 task for 325 total sessions. Animals completed 121-830 trials per session, with a median of 380. Block lengths were drawn from a uniform distribution that spanned a maximum range of 40-100 trials. For full details, we refer readers to Bari et al. (2019).

In the human dynamic foraging task, subjects performed a similar task with 4 possible states: {0.257/0.043, 0.225/0.075, 0.075/0.225, 0.043/0.257}. This dataset included 26 healthy young subjects, 26 healthy elderly control subjects, and 26 patients with Parkinson’s disease who performed the task both off and on dopaminergic medications (order counterbalanced across patients). During “off medication” sessions, patients withheld taking all dopaminergic medications for at least 10 hours. During “on medication” sessions, patients were tested an average 1.6 hours after receiving dopaminergic medications. All subjects were prescribed l-DOPA and the majority (n=17) were also taking a D_2_ receptor agonist (pramipexole, pergolide, or ropinirole). Subjects performed 800 trials in 10 blocks of 70-90 trials. We excluded one patient with Parkinson’s disease who did not complete all trials on an “on medication” session. For full details, we refer readers to Rutledge et al. (2009).

### Theoretical framework

We model an agent that can take actions (denoted by *a*) and visit states (denoted by *s*). Agents learn a policy *π*(*a*|*s*), a probabilistic mapping from states to actions. Technically, states are defined as a representation of the information needed to predict reward (Sutton and Barto, 2018). Based on this definition, the correct state for the dynamic foraging task would need to include more information than what we have included in our specification of task states given earlier. Specifically, task states correspond to the different baiting probabilities that appear repeatedly in the task, switching after a random number of trials. Because the task state is not directly observable by the agent, the state representation would need to include the sufficient statistics for the posterior probability distribution over the task state. In addition, it would need to include *t*_*a*_, the number of consecutive choices since option *a* was last chosen. These requirements significantly complicate the analysis of the optimal policy; moreover, it is doubtful that mice and humans keep accurate track of all this information at the same time.

Our model is predicated on the assumption that subjects represent a simpler state representation consisting only of the task state. Even though the task state is in fact unobservable, we restrict our analysis to behavior during “steady state” (after the first 20 trials post-switch), during which time it is plausible that subjects have relatively little task state uncertainty. The assumption that subjects neglect *t*_*a*_ (i.e., that they either do not track their choice history or do not use it in their state representation) is more drastic, but it is nevertheless common in many models of matching, and has some empirical support (Heyman, 1979; Nevin, 1969, 1979), though the evidence is equivocal (see Shimp, 1966; Silberberg et al., 1978). Further evidence against response counting is discussed later.

Policy complexity is the mutual information between states and actions:

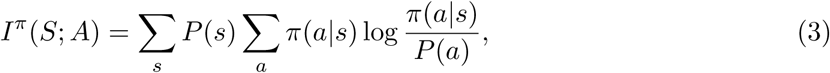

where *P* (*a*) = Σ_*s*_ *P* (*s*)*π*(*a*|*s*) is the marginal action probability. A key assumption underlying our formulation of the optimal policy is that agents are capacity constrained (i.e., there is an upper bound *C* on policy complexity). Stated another way, agents must compress the optimal policy if they lack the memory resources to store it. We therefore define a joint optimization problem, where agents seeks to maximize reward subject to a capacity constraint. We define the optimal policy as:

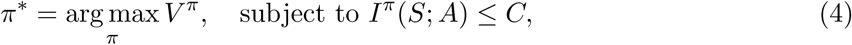

where *V* ^*π*^ is the value (average reward) under policy *π*. Note that we allow the agent to discard unnecessary information to maximize reward, so more information can never corrupt performance. The value is given by:

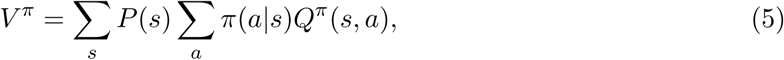

where *Q*^*π*^(*s, a*) is the average reward for taking action *a* in state *s*.

For the dynamic foraging task, the expected reward for choosing action *a* is obtained by marginalizing over *t*_*a*_:

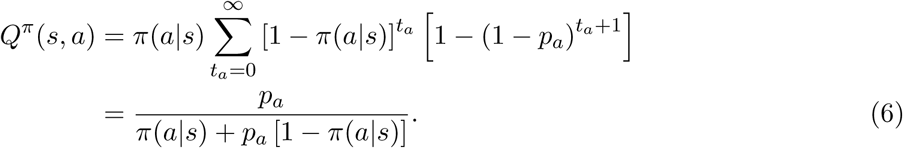

We will sometimes use the shorthand *r*_*a*_ to denote the expected reward for choosing action *a*. Because task states and actions are both marginally equiprobable, we assume in our analyses that *P* (*s*) = 1*/N* (where *N* is the number of task states) and *P* (*a*) = 1/2.

### Data analysis

Following convention, we defined undermatching by fitting the following function

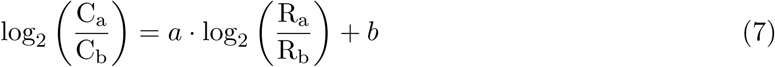

where C_a_, C_b_, R_a_, and R_b_ correspond to the total choices and rewards in an individual block. We report *a*, where *a* = 1 is perfect matching, *a* < 1 is undermatching, and *a >* 1 is overmatching (Baum, 1974). In calculating 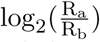, we excluded blocks with R_a_ = 0 or R_b_ = 0. For both mouse and human datasets, we excluded trials 1-20 following each block transition to allow behavior to stabilize.

To construct the empirical reward-complexity curves, in both data sets, we computed the average reward and mutual information between states and actions for each session. We estimated mutual information by computing the empirical action frequencies for each state for each session. Although there are numerous methods for computing mutual information, we found that using the empirical action frequencies for each state gave reasonably good performance, likely given the large number of trials, limited states, and limited number of actions in each state.

In performing all paired and unpaired statistical tests, we first performed a Lilliefors test, which tests the null hypothesis that the data are normally distributed. In all cases, the null hypothesis was rejected, and we subsequently performed non-parameteric testing, either the Wilcoxon rank sum test (for independent samples) or the Wilcoxon signed-rank test (for paired samples).

All code and data to reproduce the analyses in this paper can be obtained at https://github.com/bilalbari3/undermatching_compression.

## Results

### Matching is an optimal probabilistic policy in variable-interval / variable-interval tasks

To understand how policy compression leads to undermatching, we must first understand matching behavior, the task conditions that generate matching behavior, and what matching implies about the brain’s state representations. We have developed a number of these arguments previously and repeat them here for clarity (Bari and Cohen, 2021).

Task conditions are critical for observing matching behavior. Most studies employ ‘variable interval’ (VI) reward schedules. In these tasks, reward is available at an option after a variable number of choices has been made (in discrete choice tasks). Once the requisite number of choices has been made, that option is ‘baited,’ guaranteeing reward delivery when it is next chosen. The reward is not physically present, but will be delivered when that option is next chosen. The baiting rule takes the form shown in Equation 2. To gain an intuition, if the probability of reward on the unchosen option increases the longer it has been left unchosen, it makes sense to occasionally probe it to harvest baited rewards. The logic of this task rule is to mimic harvesting conditions where abstaining from a resource allows that resource to replenish. Concretely, imagine *p*_*a*_ = 0.1, *p*_*b*_ = 0.4 and the animal repeatedly chooses option *b*. On the first trial, *P*_*a*_ = 0.1 and *P*_*b*_ = 0.4. After option *b* is chosen once, *P*_*a*_ ≈ 0.19. After option *b* is chosen twice in a row, now *P*_*a*_ ≈ 0.27. As this continues, *P*_*a*_ approaches 1. After option *a* is chosen, *P*_*a*_ then resets to 0.1 and *P*_*b*_ = 0.64 since it has not been chosen for 1 trial. Figure 1A demonstrates the baiting rule for different values of *p*_*a*_.

In tasks where both options follow VI reward schedules (so-called VI/VI tasks), matching is the optimal probabilistic policy for state representations that ignore *t*_*a*_ (equation 2). Rewriting equation 1, matching occurs when 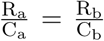. This simply states that matching occurs when the expected reward obtained from each option is equal. Figure 1B provides a geometric intuition for matching, which occurs when *r*_*a*_ and *r*_*b*_ cross one another (in this case, when *π*_*a*_ ≈ 0.86). A normative explanation is therefore that matching behavior is the best probabilistic behavior animals can exhibit to harvest reward.

### Matching is generally a suboptimal probabilistic policy and implies animals are unaware of baiting

A key insight into understanding why matching behavior arises came from Sakai and Fukai (2008). In order to implement the optimal probabilistic policy, an agent needs to understand how adjusting the parameters of its policy changes behavior (a relatively easy problem) and how changing this policy modifies the environment (a much harder problem). If an agent ignores the change in the environment, then matching behavior emerges. In other words, matching occurs because agents ignore the baiting rule (i.e., behave as if their actions do not change reward probabilities). Because of this, generally speaking, matching is not the optimal probabilistic policy.

In VI/VI tasks, matching fortuitously corresponds to the optimal probabilistic policy and we therefore cannot use this task to conclude whether agents are aware of the baiting rule. The critical test is an experiment in which matching is not the optimal strategy. For example, if one option follows a VI schedule (i.e., programmed with the baiting rule) the other follows a variable-ratio (VR) schedule (i.e., standard probabilistic reward delivery), then matching will harvest suboptimal rewards, as shown in Figure 1C.

Figure 2 demonstrates a telling set of experiments from Williams (1985), reanalyzed here. Rats performed 6 different VI/VR task variants, each summarized with one line (3 on the left plot, 3 on the right plot). Each black line here is *V* ^*π*^ for each task. In each task variant, rats more closely approximate the matching solution than the maximizing solution, and in fact demonstrate a significant degree of undermatching (choice probabilities are closer to 50% than would be expected from perfect matching). The finding that animals match instead of maximize has been replicated numerous times (Herrnstein and Heyman, 1979; Mazur, 1981; Vyse and Belke, 1992), including in humans (Savastano and Fantino, 1994).

**Figure 2:**
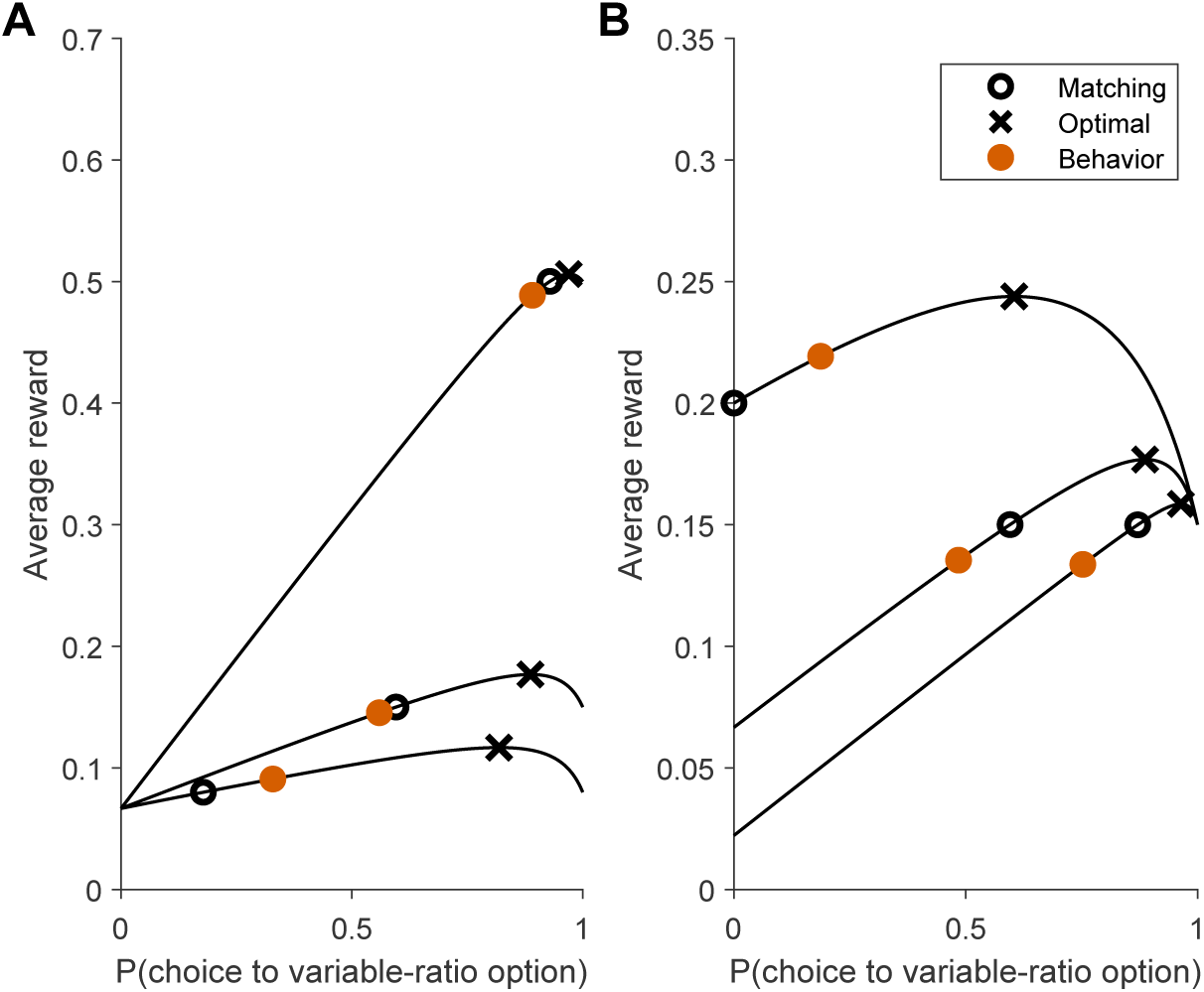
Rats are systematically biased towards undermatching instead of the optimal probabilistic policy. A: In Williams (1985), rats performed 6 different variable-interval / variable-ratio (VI/VR) task variants. Each black line demonstrates *V* ^*π*^ for each task variant. Overlaid on each *V* ^*π*^ line are **X** symbols for the optimal solution, **O** symbols for the matching solution, and filled orange circles for the empirical behavior. Rats were consistently closer to matching than to maximizing, and demonstrated a significant degree of undermatching. Because a choice of the VI option resulted in a 6s timeout, we approximated the VI option’s reward probability by 6*/τ*, where *τ* is the mean reward time under the VI schedule. The schedules are defined as follows, where each number corresponds to *p*_*i*_, the base reward probability. VI/VR, from top to bottom: (0.07, 0.5), (0.07, 0.15), (0.07, 0.08) B: VI/VR, from top to bottom: (0.2, 0.15), (0.07, 0.15), (0.02, 0.15).

We briefly note that periodic switching policies (i.e., sample the other option every *n* choices) are the global optimal policies in VI/VI tasks. These are more difficult for an agent to implement, as it requires tracking choice history (i.e., tracking *t*_*a*_ in equation 2), which necessitates a much larger state representation. In the case of *p*_*a*_ = 0.4 and *p*_*b*_ = 0.1, the optimal policy is to alternate selecting option *a* six times and option *b* once (when *P*_*b*_ ≈ 0.52). If these policies are used, they should be easy to diagnose, since stay duration distributions will be bimodal. Across a variety of species, stay duration distributions remain unimodal (Gallistel et al., 2001; Sugrue et al., 2004; Bari et al., 2019). This constitutes further evidence against models based on response counting (as discussed earlier).

Empirically, the finding that animals behave as if they are unaware of changes in environmental statistics may explain why most models of matching behavior do not account for the baiting rule. Examples include melioration (Herrnstein and Vaughan, 1980), local matching (Sugrue et al., 2004), logistic regression (Lau and Glimcher, 2005; Tsutsui et al., 2016), and models with covariance-based update rules like those underlying direct actor and actor critic agents (Loewenstein and Seung, 2006). We are aware of one study that models the baiting rule (Huh et al., 2009), though this model failed to explain behavioral data better than *Q*-learning in any of 31 mice in Bari et al. (2019).

In summary, due to the complexity of the baiting rules underlying VI/VI tasks, animals behave as if they are unaware of the baiting rule because: (1) they do not adopt the optimal probabilistic policies in VI/VR tasks; (2) they do not adopt deterministic policies in VI/VI tasks; (3) their behavior is better fit by models that ignore the baiting rule; and (4) successful models of matching typically ignore the baiting rule. Instead, animals seem to behave in these tasks as if their actions do not change the reward probabilities.

### Policy compression only allows for perfect matching or undermatching and excludes overmatching

We now apply the notion of policy compression to matching behavior. Figure 3 shows the reward-complexity frontier for the 40/10 task variant in Bari et al. (2019). In this task, mice chose between two options, each following a variable-interval schedule. One option gave reward with a base probability of *p*_*a*_ = 0.4 and the other option gave reward with *p*_*b*_ = 0.1. These alternated every 40 − 100 trials and the transitions were uncued to the mice. Using the arguments we developed above, we assume the brain believes the world consists of the following two states, (*s*_1_: *p*_*a*_ = 0.4, *p*_*b*_ = 0.1) or (*s*_2_: *p*_*a*_ = 0.1, *p*_*b*_ = 0.4). That is, we ignore the baiting rule in generating these state representations.

**Figure 3:**
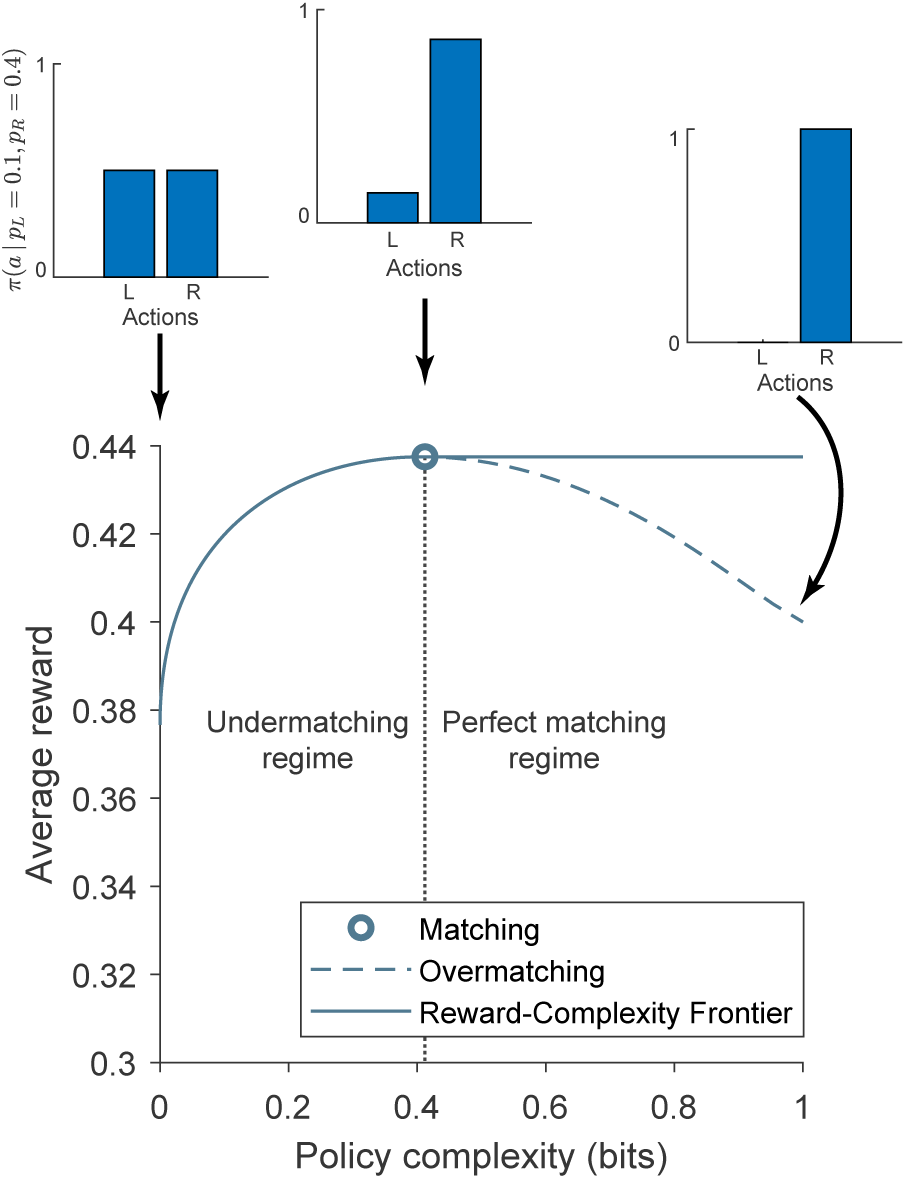
Policy compression allows for perfect matching or undermatching, but not overmatching. Under the assumption that agents are unaware of the baiting rule, the optimal reward-complexity curve is a monotonically non-decreasing function. At policy complexities below perfect matching, agents exhibit undermatching, biasing choice the poorer option (see inset for policy at policy complexity = 0 bits). At an intermediate policy complexity, agents exhibit perfect matching, harvesting optimal reward. At high levels of policy complexity, agents continue to exhibit perfect matching. Agents with sufficiently high capacity above matching are capable of exhibiting overmatching, shown by the dashed line (see inset at policy complexity = 1 bit). However, agents with sufficiently high capacity can compress their policies and increase their reward rates by adopting a perfect matching policy. Therefore, policy compression disallows overmatching.

Under this assumption, the reward-complexity frontier is a monotonically non-decreasing function. Each point on the frontier corresponds to a reward-maximizing policy under a particular policy complexity constraint. The frontier achieves a maximum at policy complexity equal to and exceeding the matching solution (Figure 3). Lower complexity policies correspond to undermatching (choosing the poorer option more often than prescribed by matching). More complex policies correspond to overmatching (choosing the richer option more often). However, overmatching yields less reward than matching with a higher cognitive cost. We posit that the brain optimizes its policy under a complexity constraint, thereby generating matching behavior.

### Mice operate near the optimal reward-complexity frontier and undermatch in a manner predicted by policy compression

Applying the policy compression framework to mouse behavior, as expected, we find that mice are capacity-constrained (Figure 4A). Moreover, they operate near the optimal frontier, which suggests that they are close to optimally balancing reward and complexity. We additionally find that policy complexity does not appreciably change across task variants (Figure 4B), suggesting a constant resource constraint, which we have observed in prior applications of policy complexity (Gershman and Lai, 2021).

**Figure 4:**
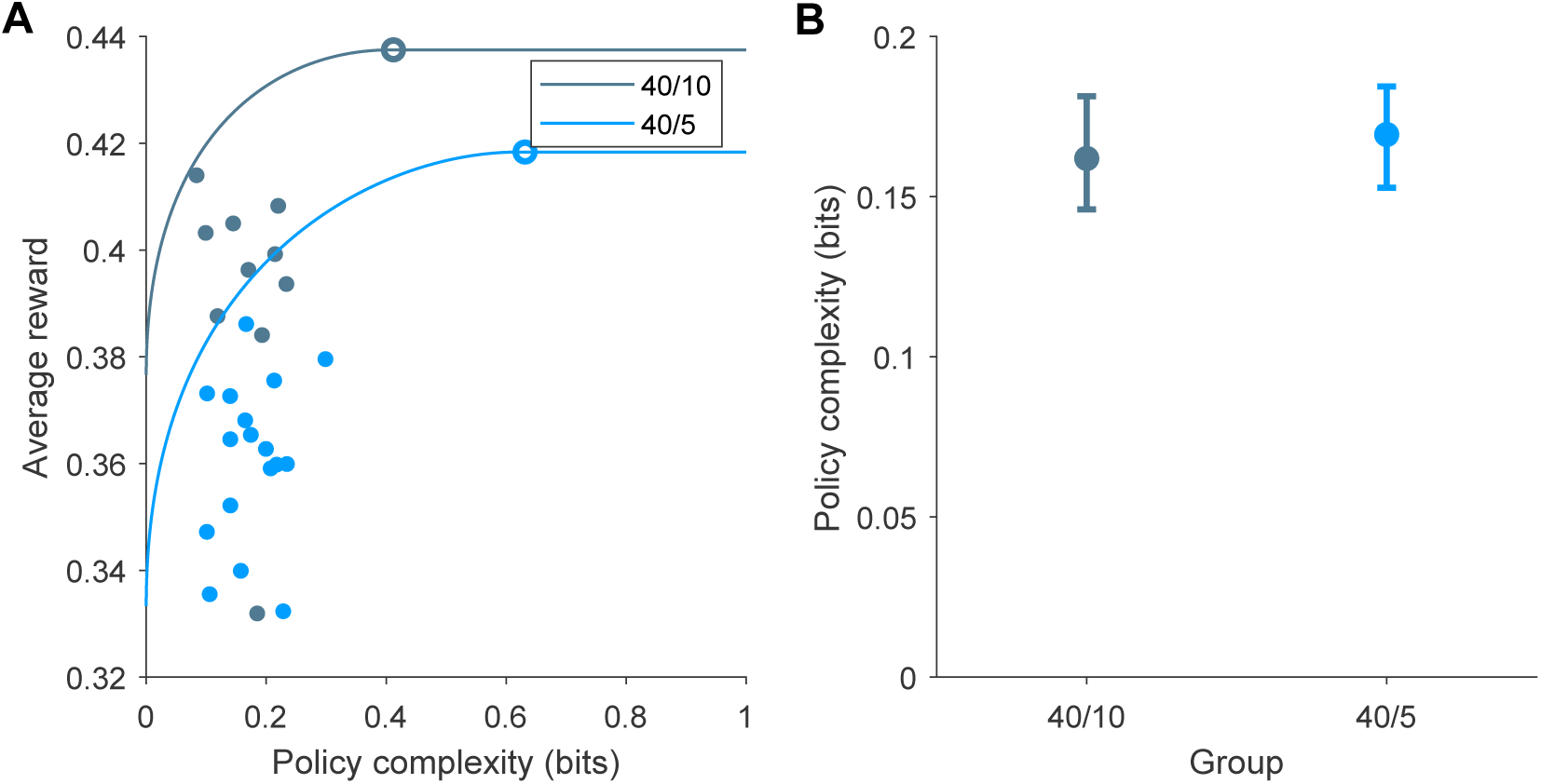
Mice exhibit capacity-constrained policies near the optimal reward-complexity frontier. A. Optimal and empirical reward-complexity curves for mice performing the 40/10 and 40/5 task variants in Bari et al. (2019). In each task variant, mice are capacity-constrained (policy complexity below optimal) and operate near the optimal reward-complexity frontier. Open circles denotes matching behavior. Filled circles denote individual mice. B. Policy complexity in each task condition. Data are median ± 95% bootstrapped CI. Wilcoxon rank-sum test, *p* = 0.18.

Policy compression makes a prediction about the degree of undermatching capacity-constrained animals should exhibit in different task variants. Undermatching should become exaggerated under reward schedules that demand more complex policies for matching behavior (Figure 5A, B). In Figure 4A, the 40/10 reward schedule demands the fewest bits for matching behavior and the 40/5 reward schedule demands more. As an alternative prediction, one might instead predict greater *overmatching* for the 40/5 reward schedule, as the higher probability side is easier to discriminate and mice may therefore perseverate on that side. We find that the policy compression prediction is borne out, with mice exhibiting significantly greater undermatching on the 40/5 schedule relative to the 40/10 schedule (Figure 5C, D).

**Figure 5:**
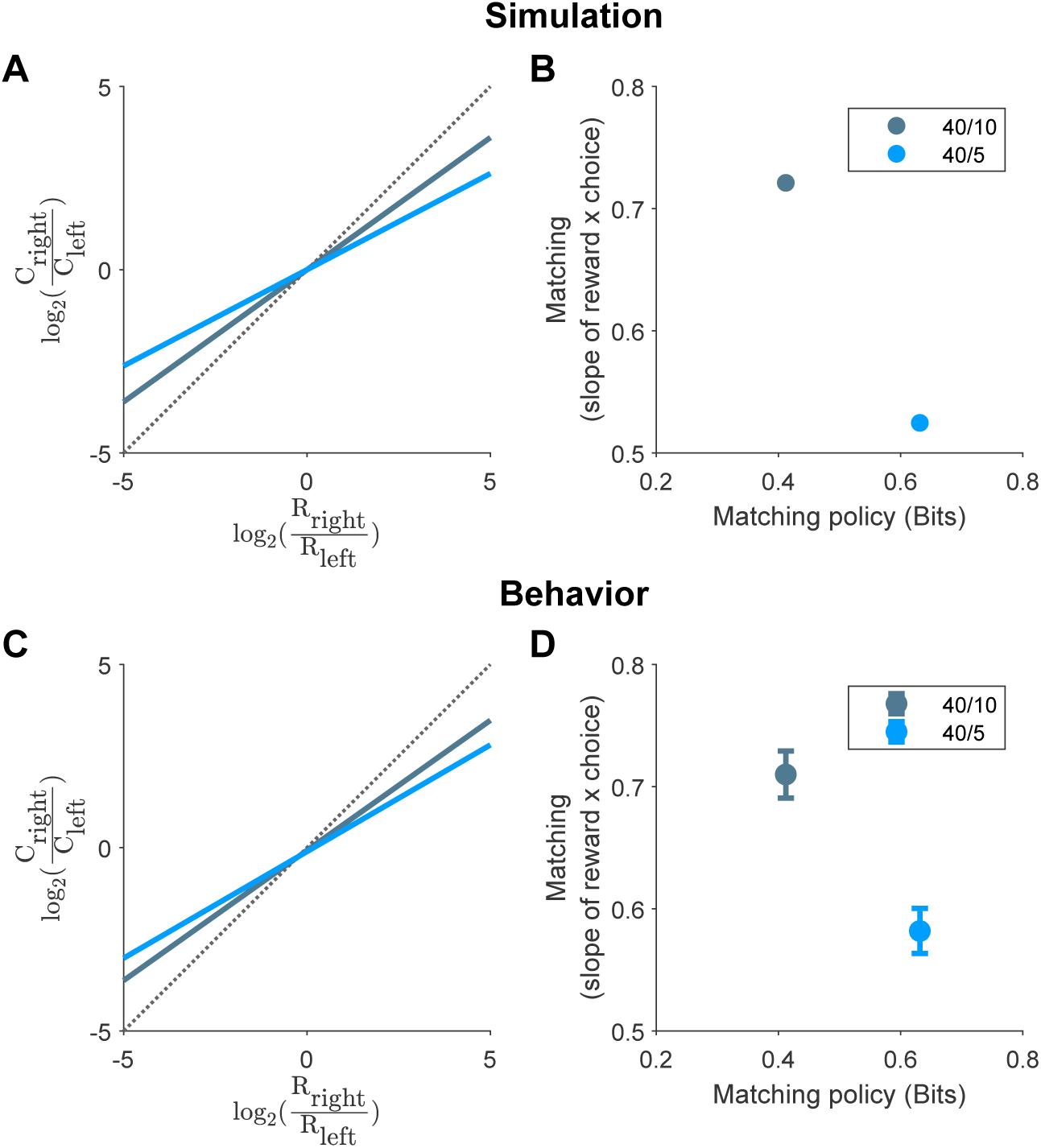
Mice exhibit greater undermatching in task variants that demand greater policy complexity. A. Simulation of a policy at a capacity constraint of 0.19 bits. The log-choice ratio is plotted as a function of log-reward ratio. Dotted line corresponds to unity. B. Theoretical matching slopes as a function of the policy complexity for perfect matching. C. Log-choice ratio as a function of log-reward ratio for the 40/10 (slope = 0.710) and 40/5 (slope = 0.582) task variants. Each colored line is the best fit and dotted line corresponds to unity. D. Empirical matching slopes (least-squares estimate ± 95% CI) for each task variant. 95% CIs: 0.691-0.729 for 40/10 task variant, 0.563-0.600 for 40/5 task variant.

### Dopaminergic medication increases capacity limits on memory

Having determined that undermatching and capacity constraint are related, we next sought to determine the neural basis underlying this capacity constraint. We have argued previously that tonic dopamine controls the precision of state representations, with greater precision accessible at a greater cognitive cost (Mikhael et al., 2021). We therefore hypothesized that tonic dopamine may modulate policy complexity, which in matching tasks would alter the degree of undermatching. Although the literature is scarce, there is some extant data to argue that pharmacologic manipulations of dopamine have the expected effect on undermatching (Soto et al., 2014; Lie et al., 2016). To address this question, we reanalyzed data from human subjects performing a dynamic foraging task, similar to the mice (Rutledge et al., 2009). Four groups of subjects (young controls, elderly controls, patients with Parkinson’s disease off dopaminergic medications, patients with Parkinson’s disease on dopaminergic medications) performed a VI/VI task, similar to the mice above. In this task, each option followed a VI schedule, with reward probabilities alternating between four task states— *s*_1_ : {0.075, 0.225}, *s*_2_ : {0.043, 0.257}, *s*_3_ : {0.225, 0.075}, *s*_4_ : {0.257, 0.043}. Subjects performed 800 trials with uncued transitions between blocks of 70 − 90 trials.

First, we confirmed that all groups demonstrated a significant degree of undermatching in this task (Figure 6). Young controls exhibited significantly less undermatching than elderly controls (Figure 7A). Interestingly, patients with Parkinson’s disease off dopaminergic medications exhibited greater undermatching than on sessions when they received dopaminergic medications, due partly to an increase in policy complexity (Figure 7B). On a subject-by-subject basis, the patients with Parkinson’s disease who demonstrated the greatest increase in policy complexity also exhibited the least degree of undermatching (Figure 7C).

**Figure 6:**
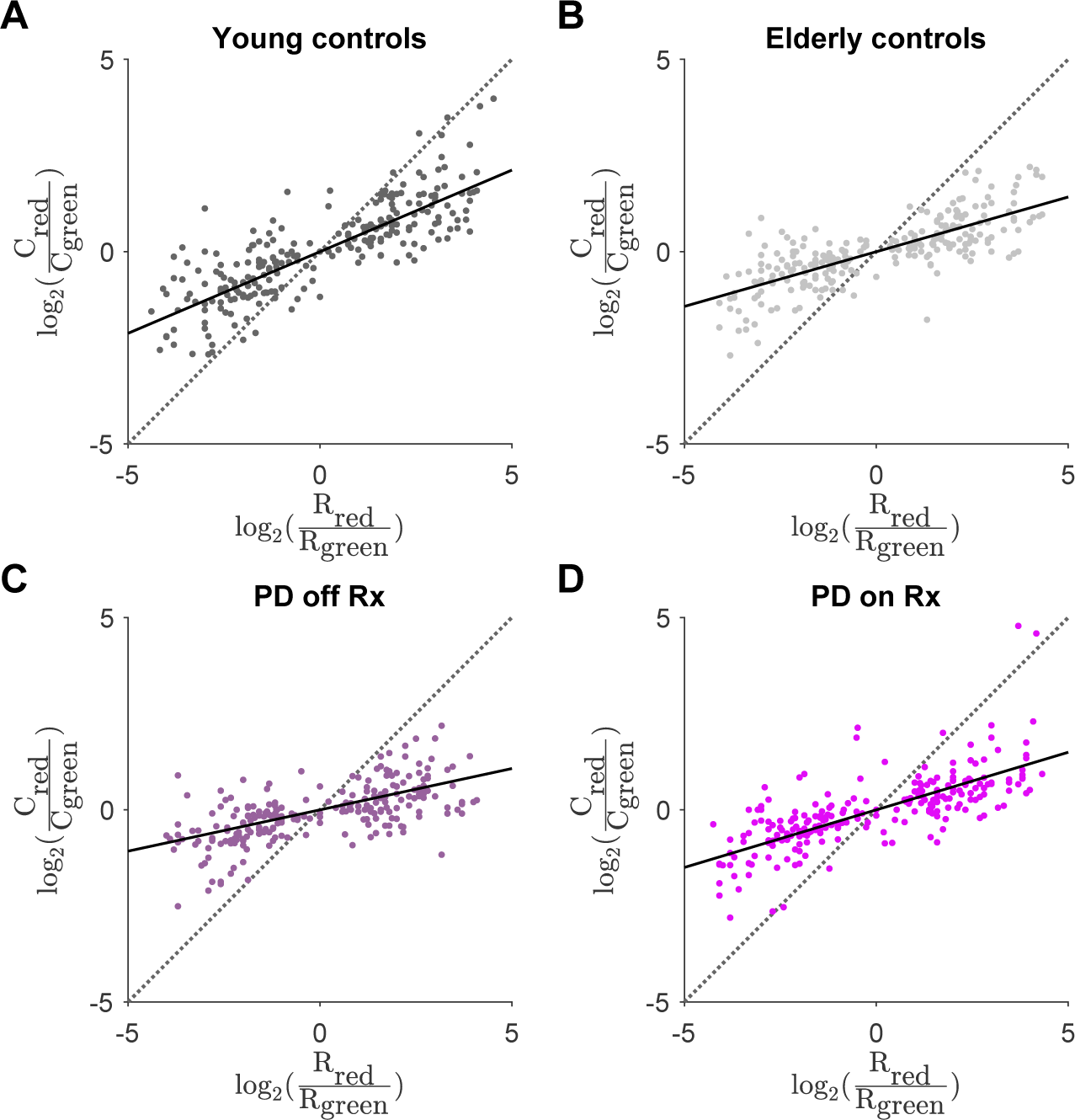
Human subjects exhibit undermatching in a dynamic foraging task. A: Log-choice ratio as a function of log-reward ratio for young control subjects. Slope = 0.424. B: Log-choice ratio as a function of log-reward ratio for elderly control subjects. Slope =0.285. C: Log-choice ratio as a function of log-reward ratio for patients with Parkinson’s disease off dopaminergic medications. Slope = 0.215. D: Log-choice ratio as a function of log-reward ratio for patients with Parkinson’s disease on dopaminergic medications. Slope = 0.300. In all panels, the black solid line corresponds to the best fit line and the dotted line corresponds to unity.

**Figure 7:**
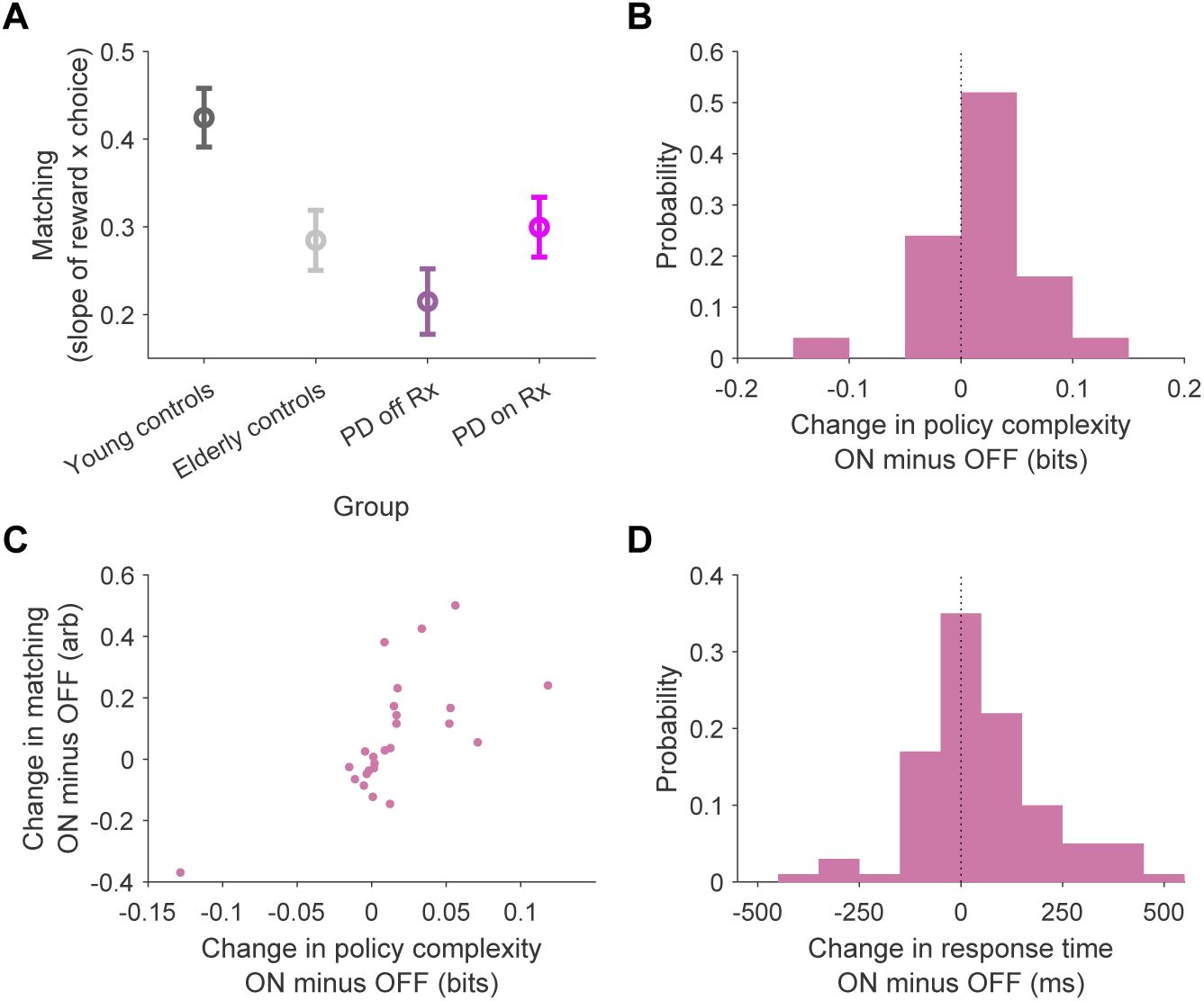
Dopaminergic medications increase policy complexity and reduce undermatching in Parkinson’s patients. A. Matching slope for each group [least-squares estimate ± 95% CI]. Young controls: 0.424 [0.391 0.458], elderly controls: 0.285 [0.250 0.319], patients with Parkinson’s disease off dopaminergic medications: 0.215 [0.178 0.252], patients with Parkinson’s disease on dopaminergic medications: 0.300 [0.265 0.334]. B. Change in policy complexity on dopaminergic medication minus off dopaminergic medication for the Parkinson’s disease group. Medians are significantly different between groups (Wilcoxon signed-rank test, *p* < 0.05). C. Change in matching as a function of change in policy complexity for the Parkinson’s disease group. Data are reported as on medication minus off medication. An increase in matching (*y*-axis) means *less* undermatching. Slope = 3.009, *p* < 0.005. D Change in response times on dopaminergic medication minus off dopaminergic medication for the Parkinson’s disease group. Medians are significantly different between groups (Wilcoxon signedrank test, *p* < 0.01).

The policy compression framework additionally makes the prediction that more complex policies should result in slower response times, since this necessitates more bits that the brain must inspect to find a coded state (Lai and Gershman, 2021). This is a counterintuitive prediction for patients with Parkinson’s disease, since bradykinesia is a defining feature of the condition that is improved with dopaminergic medications. We find that, indeed, the policy compression prediction is borne out: dopaminergic medications slow down response times for patients with Parkinson’s disease (Figure 7D).

## Discussion

Decades of empirical observations in the matching literature demonstrate a consistent bias towards undermatching, but the origin of this bias has been mysterious. Our main contribution is to show that undermatching can arise from policy compression: under some assumptions about state representation, maximizing reward subject to an upper bound on policy complexity implies undermatching or perfect matching, never overmatching. Our theory additionally makes the novel prediction that capacity-constrained agents should undermatch more on task variants that require greater policy complexity for perfect matching behavior, which we confirmed using analyses of existing data. Finally, we showed that dopaminergic medications reduce undermatching in patients with Parkinson’s disease, in part by increasing policy complexity.

We have argued that overmatching should never be observed for an agent maximizing reward under a capacity constraint. Overmatching, however, has been obtained in task variants that impose a strong cost for switching actions (Aparicio, 2001). These task variants do not pose any substantial problems for our theory. In particular, they alter both the state representation agents must use, as agents must store the last action to determine whether switching is warranted, and the calculation of expected reward *V* ^*π*^, which should include the cost of switching. Overmatching in these task variants may only appear as overmatching because these factors are excluded from analysis. We leave a more systematic treatment of this hypothesis to future work.

Our formulation of the optimal reward-complexity curve assumes lack of knowledge of the baiting rule in order to define the state representations used by agents. This yields a monotonic non-decreasing reward-complexity function (Figure 3). With a more complete state definition that includes consecutive choices of an option (Figure 1A), the reward-complexity curve should be monotonically increasing, and the policy with the greatest mutual information between states and actions would yield the greatest reward. In fact, baiting can be learned, but requires explicit training (Tunney and Shanks, 2002), and to our knowledge has not been observed simply with lengthy practice. While biological agents clearly do not possess the full state representation necessary to optimize reward in matching tasks, overtrained agents often alternate choices somewhat, which requires a representation of past choice (Lau and Glimcher, 2005; Bari et al., 2019). On occasion, this tendency to alternate can be extreme enough to warrant a ‘changeover delay,’ a penalty imposed on switching (Herrnstein, 1961; Sugrue et al., 2004), although this does not often need to be implemented. Although alternation appears to be a minor component of behavior, it is one that emerges with training, and it is unclear how agents learn this augmented state representation.

We make no claims about the particular algorithms for optimizing the reward-complexity trade-off. While we know the capacity-constrained policy that would maximize reward (Gershman, 2020; Lai and Gershman, 2021), agents are often suboptimal in harvesting reward for a particular policy complexity (Figure 4A). This suboptimality persists despite analyzing only steady-state behavior, suggesting it is not only due to slow learning at block transitions. Because mice made a choice by licking a leftward or rightward lick port, rather than an abstract color-coded option in the human task, they exhibited notable side-specific biases, favoring one option over the other. The origin of this bias and why it persists despite being detrimental to reward harvesting is largely unknown.

One intriguing possibility here is that policy compression may be adaptive. We have argued that agents unaware of the baiting rule benefit from policy compression - here, by adopting perfect matching rather than overmatching when above capacity. It remains an open question whether agents with impoverished state representations may benefit from policy compression more generally, and under what conditions. Prior work has demonstrated that undermatching may increase reward received in dynamic foraging tasks, due to the nonstationary nature of the task (Iigaya et al., 2019). It has been suggested that biological agents possess an inductive bias towards environmental volatility (Yu and Cohen, 2008) and this belief has been proposed as an explanation for undermatching (Saito et al., 2014) and matching more broadly (Yu and Huang, 2014). Given the relationship between undermatching and exploration—undermatching corresponds to low capacity, which produces more random behavior (Lai and Gershman, 2021)—an alternative perspective is that agents exhibit elevated random exploration in task variants with high volatility.

## Acknowledgements

We are indebted to Robb B. Rutledge and Jeremiah Y. Cohen for making their data available.

## References

Anderson, K. G., Velkey, A. J., and Woolverton, W. L. (2002). The generalized matching law as a predictor of choice between cocaine and food in rhesus monkeys. Psychopharmacology, 163:319–326.

Aparicio, C. F. (2001). Overmatching in rats: The barrier choice paradigm. Journal of the Experimental Analysis of Behavior, 75(1):93–106.

Bari, B. A. and Cohen, J. Y. (2021). Dynamic decision making and value computations in medial frontal cortex. International Review of Neurobiology, 158:83.

Bari, B. A., Grossman, C. D., Lubin, E. E., Rajagopalan, A. E., Cressy, J. I., and Cohen, J. Y. (2019). Stable representations of decision variables for flexible behavior. Neuron, 103:922–933.

Baum, W. M. (1974). On two types of deviation from the matching law: bias and undermatching. Journal of the Experimental Analysis of Behavior, 22:231–242.

Baum, W. M. (1979). Matching, undermatching, and overmatching in studies of choice. Journal of the Experimental Analysis of Behavior, 32:269–281.

Beardsley, S. D. and McDowell, J. (1992). Application of Herrnstein’s hyperbola to time allocation of naturalistic human behavior maintained by naturalistic social reinforcement. Journal of the Experimental Analysis of Behavior, 57:177–185.

Belke, T. W. and Belliveau, J. (2001). The generalized matching law describes choice on concurrent variable-interval schedules of wheel-running reinforcement. Journal of the Experimental Analysis of Behavior, 75:299–310.

Cero, I. and Falligant, J. M. (2020). Application of the generalized matching law to chess openings: A gambit analysis. Journal of Applied Behavior Analysis, 53:835–845.

Cohen, J. Y., Haesler, S., Vong, L., Lowell, B. B., and Uchida, N. (2012). Neuron-type-specific signals for reward and punishment in the ventral tegmental area. Nature, 482:85–88.

de Villiers, P. A. and Herrnstein, R. J. (1976). Toward a law of response strength. Psychological Bulletin, 83:1131–1153.

Fonseca, M. S., Murakami, M., and Mainen, Z. F. (2015). Activation of dorsal raphe serotonergic neurons promotes waiting but is not reinforcing. Current Biology, 25:306–315.

Gallistel, C. (1994). Foraging for brain stimulation: toward a neurobiology of computation. Cognition, 50:151–170.

Gallistel, C., King, A. P., Gottlieb, D., Balci, F., Papachristos, E. B., Szalecki, M., and Carbone, K. S. (2007). Is matching innate? Journal of the Experimental Analysis of Behavior, 87:161–199.

Gallistel, C., Mark, T. A., King, A. P., and Latham, P. (2001). The rat approximates an ideal detector of changes in rates of reward: implications for the law of effect. Journal of Experimental Psychology: Animal Behavior Processes, 27:354—372.

Gershman, S. J. (2020). Origin of perseveration in the trade-off between reward and complexity. Cognition, 204:104394.

Gershman, S. J. and Lai, L. (2021). The reward-complexity trade-off in schizophrenia. Computational Psychiatry, 5.

Graft, D., Lea, S., and Whitworth, T. (1977). The matching law in and within groups of rats. Journal of the Experimental Analysis of Behavior, 27:183–194.

Herrnstein, R. J. (1961). Relative and absolute strength of response as a function of frequency of reinforcement. Journal of the Experimental Analysis of Behavior, 4:267–272.

Herrnstein, R. J. and Heyman, G. M. (1979). Is matching compatible with reinforcement maximization on concurrent variable interval, variable ratio? Journal of the Experimental Analysis of Behavior, 31:209–223.

Herrnstein, R. J. and Vaughan, W. (1980). Melioration and behavioral allocation. Limits to action: The allocation of individual behavior, pages 143–176.

Heyman, G. (1979). A Markov model description of changeover probabilities on concurrent variable-interval schedules. Journal of the Experimental Analysis of Behavior, 31:41–51.

Huh, N., Jo, S., Kim, H., Sul, J. H., and Jung, M. W. (2009). Model-based reinforcement learning under concurrent schedules of reinforcement in rodents. Learning & Memory, 16:315–323.

Iigaya, K., Ahmadian, Y., Sugrue, L. P., Corrado, G. S., Loewenstein, Y., Newsome, W. T., and Fusi, S. (2019). Deviation from the matching law reflects an optimal strategy involving learning over multiple timescales. Nature Communications, 10:1–14.

Iigaya, K. and Fusi, S. (2013). Dynamical regimes in neural network models of matching behavior. Neural Computation, 25:3093–3112.

Kubanek, J. and Snyder, L. H. (2015). Matching behavior as a tradeoff between reward maximization and demands on neural computation. F1000Research, 4.

Lai, L. and Gershman, S. J. (2021). Policy compression: An information bottleneck in action selection. In Psychology of Learning and Motivation, volume 74, pages 195–232. Elsevier.

Lau, B. and Glimcher, P. W. (2005). Dynamic response-by-response models of matching behavior in rhesus monkeys. Journal of the Experimental Analysis of Behavior, 84:555–579.

Lee, S.-H., Huh, N., Lee, J. W., Ghim, J.-W., Lee, I., and Jung, M. W. (2017). Neural signals related to outcome evaluation are stronger in CA1 than CA3. Frontiers in Neural Circuits, 11:40.

Lie, C., Macaskill, A. C., and Harper, D. N. (2016). The effect of MDMA on sensitivity to rein-forcement rate. Behavioral Neuroscience, 130:243.

Loewenstein, Y. (2008). Robustness of learning that is based on covariance-driven synaptic plasticity. PLoS Computational Biology, 4:e1000007.

Loewenstein, Y. and Seung, H. S. (2006). Operant matching is a generic outcome of synaptic plasticity based on the covariance between reward and neural activity. Proceedings of the National Academy of Sciences, 103:15224–15229.

Mazur, J. E. (1981). Optimization theory fails to predict performance of pigeons in a two-response situation. Science, 214:823–825.

Mikhael, J. G., Lai, L., and Gershman, S. J. (2021). Rational inattention and tonic dopamine. PLoS Computational Biology, 17:e1008659.

Myers, D. L. and Myers, L. E. (1977). Undermatching: A reappraisal of performance on concurrent variable-interval schedules of reinforcement. Journal of the Experimental Analysis of Behavior, 27:203–214.

Nevin, J. (1969). Interval reinforcement of choice behavior in discrete trials. Journal of the Experimental Analysis of Behavior, 12:875–885.

Nevin, J. A. (1979). Overall matching versus momentary maximizing: Nevin (1969) revisited. Journal of Experimental Psychology: Animal Behavior Processes, 5:300–306.

Niv, Y., Daw, N. D., Joel, D., and Dayan, P. (2007). Tonic dopamine: opportunity costs and the control of response vigor. Psychopharmacology, 191:507–520.

Parush, N., Tishby, N., and Bergman, H. (2011). Dopaminergic balance between reward maximization and policy complexity. Frontiers in Systems Neuroscience, 5:22.

Pierce, W. D. and Epling, W. F. (1983). Choice, matching, and human behavior: A review of the literature. The Behavior Analyst, 6:57–76.

Rutledge, R. B., Lazzaro, S. C., Lau, B., Myers, C. E., Gluck, M. A., and Glimcher, P. W. (2009). Dopaminergic drugs modulate learning rates and perseveration in Parkinson’s patients in a dynamic foraging task. Journal of Neuroscience, 29:15104–15114.

Saito, H., Katahira, K., Okanoya, K., and Okada, M. (2014). Bayesian deterministic decision making: a normative account of the operant matching law and heavy-tailed reward history dependency of choices. Frontiers in Computational Neuroscience, 8:18.

Sakai, Y. and Fukai, T. (2008). When does reward maximization lead to matching law? PLoS One, 3:e3795.

Savastano, H. I. and Fantino, E. (1994). Human choice in concurrent ratio-interval schedules of reinforcement. Journal of the Experimental Analysis of Behavior, 61:453–463.

Schroeder, S. R. and Holland, J. G. (1969). Reinforcement of eye movement with concurrent schedules. Journal of the Experimental Analysis of Behavior, 12:897–903.

Shimp, C. (1966). Probabilistically reinforced choice behavior in pigeons. Journal of the Experimental Analysis of Behavior, 9:443–455.

Silberberg, A., Hamilton, B., Ziriax, J. M., and Casey, J. (1978). The structure of choice. Journal of Experimental Psychology, 4:368–398.

Soltani, A., Lee, D., and Wang, X.-J. (2006). Neural mechanism for stochastic behaviour during a competitive game. Neural Networks, 19:1075–1090.

Soltani, A., Rakhshan, M., Schafer, R. J., Burrows, B. E., and Moore, T. (2021). Separable influences of reward on visual processing and choice. Journal of Cognitive Neuroscience, 33:248–262.

Soto, P. L., Hiranita, T., Grandy, D. K., and Katz, J. L. (2014). Choice for response alternatives differing in reinforcement frequency in dopamine d2 receptor mutant and swiss-webster mice. Psychopharmacology, 231:3169–3177.

Sugrue, L. P., Corrado, G. S., and Newsome, W. T. (2004). Matching behavior and the representation of value in the parietal cortex. Science, 304:1782–1787.

Sutton, R. S. and Barto, A. G. (2018). Reinforcement Learning: An Introduction. MIT Press.

Trepka, E., Spitmaan, M., Bari, B. A., Costa, V. D., Cohen, J. Y., and Soltani, A. (2021). Entropy-based metrics for predicting choice behavior based on local response to reward. Nature Communications, 12:1–16.

Tsutsui, K.-I., Grabenhorst, F., Kobayashi, S., and Schultz, W. (2016). A dynamic code for economic object valuation in prefrontal cortex neurons. Nature Communications, 7:1–16.

Tunney, R. J. and Shanks, D. R. (2002). A re-examination of melioration and rational choice. Journal of Behavioral Decision Making, 15:291–311.

Villarreal, M., Velázquez, C., Baroja, J. L., Segura, A., Bouzas, A., and Lee, M. D. (2019). Bayesian methods applied to the generalized matching law. Journal of the Experimental Analysis of Behavior, 111:252–273.

Vullings, C. and Madelain, L. (2018). Control of saccadic latency in a dynamic environment: Allocation of saccades in time follows the matching law. Journal of Neurophysiology, 119:413–421.

Vyse, S. A. and Belke, T. W. (1992). Maximizing versus matching on concurrent variable-interval schedules. Journal of the Experimental Analysis of Behavior, 58:325–334.

Wearden, J. and Burgess, I. (1982). Matching since baum (1979). Journal of the Experimental Analysis of Behavior, 38:339–348.

Williams, B. A. (1985). Choice behavior in a discrete-trial concurrent VI-VR: A test of maximizing theories of matching. Learning and Motivation, 16:423–443.

Yu, A. J. and Cohen, J. D. (2008). Sequential effects: Superstition or rational behavior? Advances in Neural Information Processing Systems, 21.

Yu, A. J. and Huang, H. (2014). Maximizing masquerading as matching in human visual search choice behavior. Decision, 1:275–287.

